# Population genomic analyses reveal extensive genomic regions within selective sweeps associated with adaptation and demographic history of a wheat fungal pathogen

**DOI:** 10.1101/2023.08.29.555254

**Authors:** Yun Xing, Chongjing Xia, Liang Huang, Na Zhao, Hongfu Li, Xingzong Zhang, AGe Qiu, Wanqiang Tang, Meinan Wang, Xianming Chen, Bo Liu, Hao Zhang, Li Gao, Wanquan Chen, Taiguo Liu

## Abstract

Plant pathogens can quickly evolve in response to the changing host and environment, resulting in destructive epidemics. The fungus *Puccinia striiformis* f. sp. *tritici* (*Pst*), causing wheat stripe rust disease worldwide, is one such pathogen. In the arms race between pathogens and hosts, changes in host populations exerted continuous selective forces on the dynamics of pathogens. However, the footprints of these selection forces and the demography of *Pst* remain poorly explored, limited our understanding the evolutionary processes that shape the spread and evolution of pathogens. In this study, we revealed several features of worldwide *Pst* populations through population genomic analyses. There might be limited gene flow between *Pst* populations from China and other countries. A slower rate of linkage disequilibrium decay was detected comparing to rust fungi with known sexual reproduction. Furthermore, we detected extensive hard and soft sweeps associated with *Pst* adaptation. Genes within the selective sweeps were enriched in secreted proteins and effectors and showed functions related to pathogenicity or virulence, temperature tolerance, and fungicide resistance implying that Chinese *Pst* populations suffered positive selection pressures from host and abiotic factors. Moreover, demographic history indicated *Pst* populations experienced strong bottlenecks at the beginning of the wheat domestication around 10,000 years ago and during modern agriculture around 100 years ago, suggesting that the crop domestication and breeding programs could continuously contribute to the decline of pathogen effective population sizes. Our results provided insights into the evolution of the *Pst* genome and highlighted the role of modern agriculture on pathogen demography.

**Author Summary:** Wheat stripe rust, as a pathogen of wheat, is capable of inducing significant yield reduction on a global scale. Traditional epidemiological investigations have elucidated the causative agent behind this disease, the wheat stripe rust pathogen, as an organism exhibiting swift adaptive evolution in response to its environment.

However, the precise mechanisms underpinning its adaptive strategies and the ensuing alterations in population dynamics remain relatively unexplored.

In our research, we have undertaken an analysis encompassing wheat stripe rust samples throughout the world. Employing population genomics methods, we have sought to discern the genomic footprints corresponding to the adaptive process within the pathogen. These genomic footprints have illuminated the operation of positive selection pressures caused by both the host plant and the ecological context.

Concurrently, it have provided insights into the population size that the wheat stripe rust pathogen has undergone over millennia. Notably, its change in population size revealed the demographic history associated with wheat domestication.

## Introduction

Fungi are ubiquitous and play fundamental roles in ecosystems. Among them is a group of invasive fungi that cause severe diseases on plant and jeopardize the global food security (Velásquez et al., 2017; Ristaino et al., 2021). These invasive plant fungal pathogens generally have the ability to rapidly evolve in response to novel environments, particularly in the agriculture ecosystem (Croll & McDonald, 2017; Wan et al., 2007; Milus et al., 2008; Estep et al., 2015). The consequences of the rapid evolution include either the emergence of disease on new host species/cultivars, or the re-emergence of the pathogen with greater pathogenicity. Regardless which way, they become more adaptive and increase the potential of further epidemics. Understanding the biological and evolutionary features underlying the invasive plant fungal pathogens is vital for implementation of effective control strategies, and is one of the tasks in population genetics and molecular ecology (Gladieux et al., 2015).

*Puccinia striiformis* f. sp. *tritici* (*Pst*) is a wheat fungal pathogen, causing the stripe rust disease on wheat. This disease is one of the most destructive wheat diseases worldwide, especially in China, the US, and Europe (Wan et al., 2007; Chen et al., 2013). *Pst* is a heteroecious fungus, requiring two taxonomically distinct species to complete its life-cycle. The asexual stage, which is the most common stage, is on wheat (*Triticum aestivum*), producing urediniospores. Severe epidemics of wheat stripe rust are all caused by the asexual urediniospores since several cycles of asexual reproduction can be completed in a single wheat season. The sexual stage, which was unknown until recently, is on *Berberis* and *Mahonia* (Jin et al., 2010; Zhao et al., 2023). Although tens of *Berberis* and *Mahonia* species could be served as alternate hosts for *Pst,* the frequency of the sexual reproduction was low partially due to the incompatible phenology of the hosts (Zhao et al., 2016). It remains still unclear the frequency of the sexual reproduction of *Pst* nature populations. The most effective strategy for controlling this disease is the deployment of resistant wheat cultivars (Chen, 2014). However, like other invasive fungi, *Pst* could evolve rapidly, resulting in the emergence of new aggressive races (*aka* pathotypes). These new races rendered the widely grown resistant cultivars ineffective in just a few years after their release (Milus et al., 2006; Milus et al., 2008; Esmail et al., 2021; Wellings et al., 2011). In fact, several documented epidemics in the recent history were all resulted from the emergence of new *Pst* races, for example, the breakdown of the *Yr17* resistance in the 1990s in northern Europe (Bayles et al., 2000). So far, a depth understanding of the evolutionary processes underlying the rapid evolving of *Pst* is still lacking.

Survival of hash environments (e.g., the scorching summer and chilly winter) is critical for plant pathogens and the subsequent disease development. For *Pst*, oversummering and overwintering are two bottlenecks in the disease cycle. Since the wheat growing areas in China have great variations in geomorphological and climatological features, the pathogen can both oversummer and overwinter even though in different regions. Therefore, it is believed that, together with the field survey, the wheat stripe rust has an independent epidemic system in China (Zeng & Luo 2008; Chen et al., 2013). Based on geography, cropping system, extensive field survey, and long-term monitoring, the main areas where stripe rust is prevalent in China are divided into three epidemiological regions. The first region mainly covers the provinces of Yunnan, Chongqing, Guizhou, and their neighboring regions, where *Pst* can continually multiply in winter and served as sources of inoculum for local and northern regions in the following spring. So this region is called the SprS region. The second epidemiological region is called Autumn sources of inoculum (AutS), covering southern Gansu, northwest Sichuan, and the adjacent regions. The third epidemiological region named Spring epidemic areas (SprE) includes wheat-growing areas of the Huang-Huai-Hai Plain and the lower reaches of the Yangtze River (Chen et al., 2013). At the global scale, it was believed that there was little chance of population exchange between *Pst* populations in China and that in other countries because of the geographical barriers (Zeng & Luo 2006). On the other hand, the wheat stripe rust is an air-born disease and can disperse to a long distance through wind. Even though few studies have been conducted to examine the population diversity and differentiation between *Pst* populations from China and other countries, only limited number of isolates from China and limited number of molecular markers were used (Ali et al., 2014; Ding et al., 2021). The potential of admixture of *Pst* population at the global scale needs to be further evaluated.

In a successful infection, the pathogen has to invade several layers of host immune systems (Jones & Dangl 2006). In the first layer, the microbial/pathogen-associated molecular patterns (MAMPs/PAMPs) are detected by plant pattern recognition receptors, resulting in pattern-triggered immunity (PTI) in plants. The pathogens then deliver effectors (typically small secreted molecules) to suppress the PTI, resulting in the susceptibility of plants (Giraldo & Valent, 2013). In the next layer, the effector is recognized by the resistance product of plant, leading to effector-triggered immunity (ETI). Then, during the evolution, the pathogens equipped with new effectors or lost the previous effectors will be selected in the population, so that the pathogens can overcome the plant ETI. In this arms race between pathogens and hosts during evolution, changes in host populations exerted continuous selective forces on the dynamics of pathogens (Jones & Dangl 2006; Presti et al., 2015). This co-evolution provides good opportunities to detect the footprints of pathogen adaptation in genomes (Vitti et al., 2013; Everhart et al., 2021; Badouin *et al*., 2017; Duan et al., 2010; Hartmann et al., 2018; Mohd-Assaad et al., 2018; Grandaubert et al., 2019; Miller et al., 2020; Schweizer et al., 2021). Specifically in the genome of *Pst*, hundreds of effector-coding genes have been annotated, but the functions are not known for most of them. Therefore, the detection of candidate effectors under selection will be valuable for both function studies and the understanding the rapid adaptation of *Pst*.

In the pathogen-host interactions, it still remains unclear that how the domestication of crops has an impact on the evolution of pathogens. Crop domestication has often resulted in a significant reduction of the genetic variation in the crop population (Milla et al., 2015; Silk & Hodgson 2020). The reduced genetic variation of crop population may cause bottleneck effects for pathogen population during crop domestication. Recently, high throughput genomic sequencing enabled the inference of the population demography in more detail (Grünwald et al., 2016; Beichman et al., 2018; Schweizer et al., 2021; Skrede et al., 2021; Kozhar et al., 2022). *Pst* is an obligate biotroph, meaning that it can only infect and obtain nutrient from the living tissue. Such an intimate association provides an opportunity to study the demography of *Pst* during the wheat domestication. This will further provide insights into the prediction of pathogen population changes when host resistance is utilized to control the disease in modern agriculture.

Here, we aimed to reveal evolutionary features that might be associated with the rapid adaptation of *Pst* pathogen on wheat. We performed whole-genome sequencing (WGS) on 69 *Pst* isolates collected from across China in 2015. Besides, we retrieved 28 published *Pst* genome sequences from the known geographical distribution, including India, Europe, the USA, Australia and eastern Africa to represent the international population for the species. We then applied population genomic analyses to answer the following specific questions: 1) what is the genetic make-up of the *Pst* populations? Does migration or admixture exist between *Pst* populations from China and other countries? 2) what is the degree of sexual reproduction for *Pst*, particularly comparing to the sister species *P. graminis* f. sp. *tritici* which had well-known sexual reproduction? 3) are there footprints of selective sweeps in the genome of *Pst*? Do these genomic regions involve effector genes that are associated with adaptation? 4) Can we infer the demographic history for *Pst*? How is that affected by the wheat domestication? This information will enable us to better understand the adaptation of this destructive pathogen and to evaluate the potential impact of modern agriculture on the evolution of pathogens.

## Results

For the 69 Chinese *Pst* isolates, between 38.16% and 78.73% of reads were mapped to the reference genome of CYR34, and the sequencing depth ranged from 24.38X to 47.07X (S1 Table). Sequencing data from 28 isolates from Australia, India, Ethiopia, Denmark, France, the UK, and the USA with mapping coverage greater than 20X were retrieved and used in subsequent analyses (S2 Table). After quality control and filtering, 783,514 bi-allelic SNPs were called from all 97 isolates for the following analyses, except in population structure analysis in which 46,152 SNPs were used after thinning by 1 kb.

### Population structure and diversity

We inferred population structure with phylogeny and ADMIXTURE using whole-genome diversity data (Fig 1). ADMIXTURE identified the most optimal cluster at K=7 (S1 Fig) and a cluster of admixture isolates was also defined when the ancestry was below 70%. Overall, the phylogenetic clades and ADMIXTURE clusters revealed the similar population structures. For instance, the phylogenetic clade I and the ADMIXTURE clusters G1 and G4 contained exclusively only the non-China isolates. In fact, almost all (26 out of the total of 28) non-China isolates were exclusively grouped to clade I without any isolates from China. We also noticed that the four non-China isolates grouped to Admixture had distinct ancestry composition patterns (containing G3, G4, and G7 ancestries) comparing with the admixture isolates from China. This was more apparent when K=8 (S1 Fig). Given that only two out of the total of 28 non-China isolates had similar ancestry composition with the isolates from China, we speculated that there might be only limited gene flow between the *Pst* isolates from China and non-China. The PCA analysis showed a similar clustering pattern in which *Pst* isolates from China were grouped into two clusters and separated from the remaining non-China isolates (Fig 2a).

**Fig 1.**
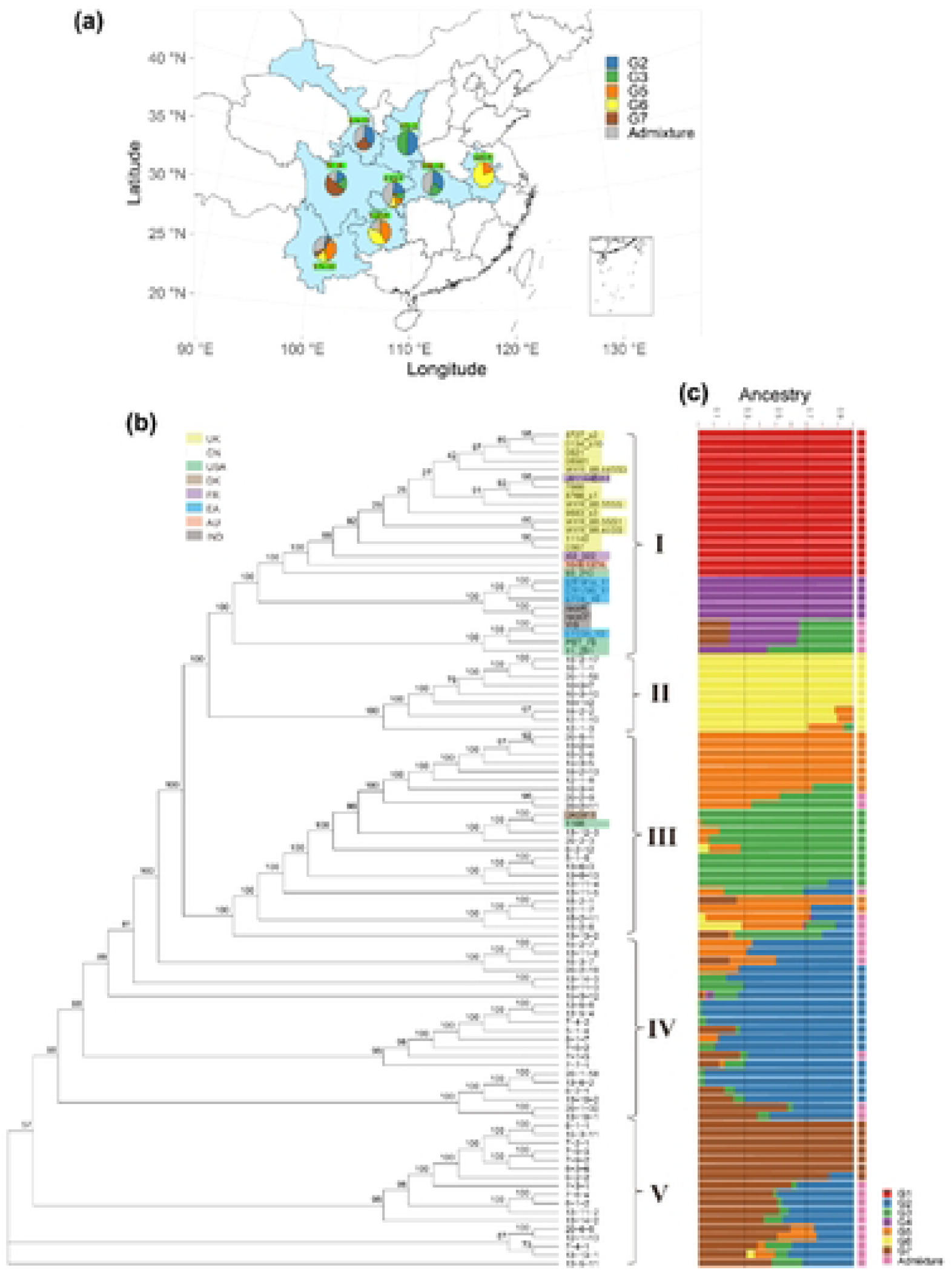
Sampling information, phylogenetic relationship, and population structure of the *Puccinia striiformis* f. sp. *tritici* (*Pst*) isolates used in this study. (a) Sampling information about provinces, number of isolates, and composition of genetic ancestries of the 69 isolates in China. (b) A phylogenetic relationship of total 98 *Pst* isolates used in this study. The neighbor-joining tree was constructed based on the genetic distance estimated using genome-wide SNPs. For visualization purpose, the tree branches were scaled to same length but not proportional to mutation rate. The numbers around the phylogenetic tree nodes were bootstrap values. (c) The population structure inferred from ADMIXTURE at the optimal number of 7 clusters. The isolates were designated to Admixture cluster when the highest ancestry to specific cluster was below 0.7. CN, China; AU, Austrilia; IND, India; EA, Eastern Africa; EU, Europe; USA, the United States of America; DK, Denmark; FR, France.

**Fig 2.**
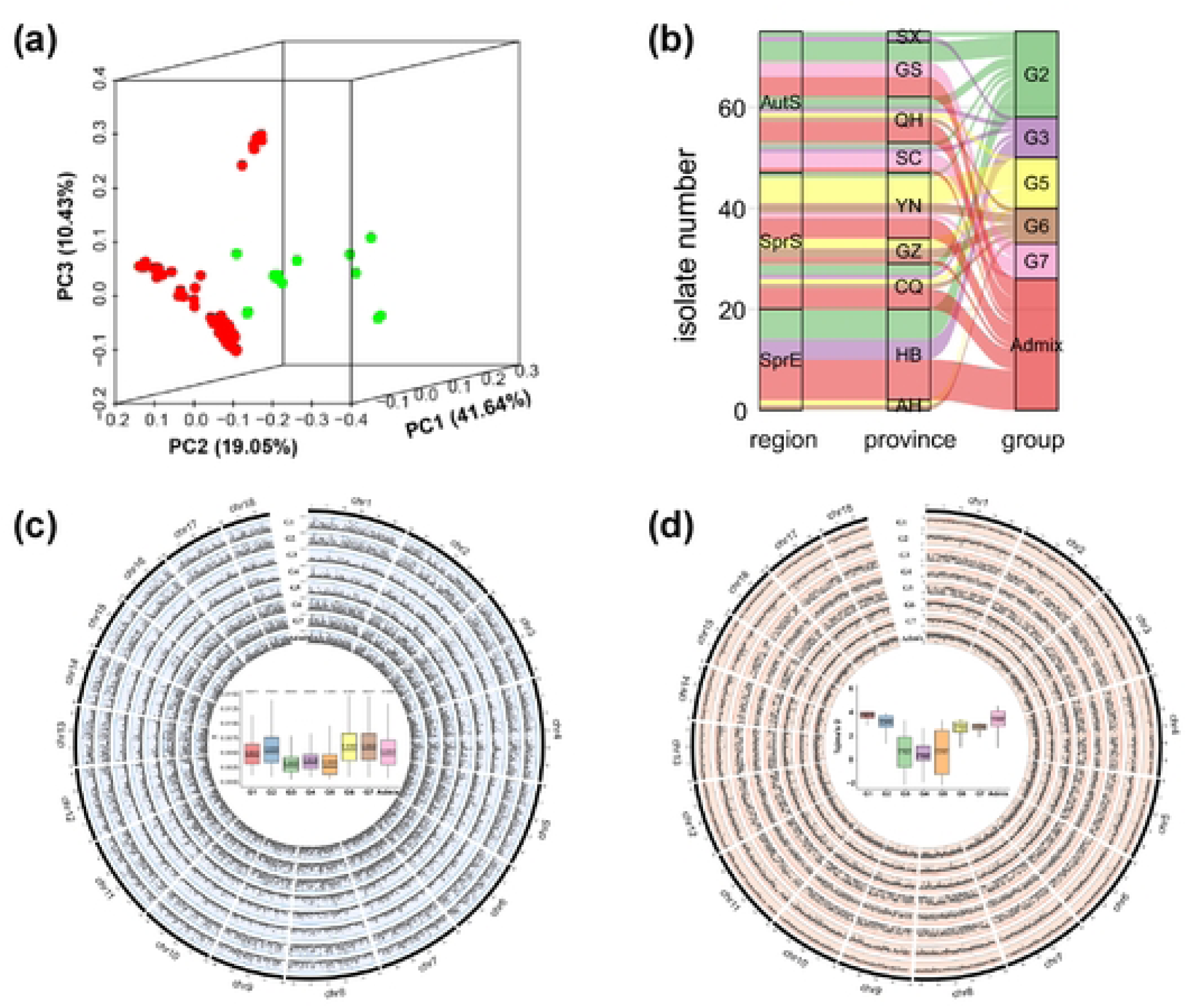
Clustering and population diversity of *Puccinia striiformis* f. sp. *tritici* isolates. (a) 3D scatter plot of the principal component analysis of 98 worldwide *Pst* isolates using whole-genome wide SNPs. The isolates from China and non-China were represented by solid red and green circles, respectively. The first three principal components explain 41.64%, 19.05%, and 10.43% of the overall variance. (b) Sankey plot of 69 *Pst* isolates from China across epidemiological regions, provinces, and genetic clusters from ADMIXTURE. AutS, Autumn sources of inoculum; SprS, Spring sources of inoculum; SprE, Spring epidemic areas; SX, Shanxi; GS, Gansu; QH, Qinghai; SC, Sichuan; YN, Yunnan; GZ, Guizhou; CQ, Chongqing; HB, Hubei; AH, Anhui. (c) nucleotide diversity (θ_π_) of ADMIXTURE-defined *Pst* populations. The circos plot shows the distribution of θ_π_ along the chromosomes for each population. The boxplot shows the summary of θ_π_for each population. Solid lines in the boxplot represent the mean, numbers above each boxplots are total number of biallelic SNPs detected in the population. (f) Tajima’s *D* of ADMIXTURE-defined *Pst* populations. Solid lines in the boxplot represent the mean, numbers above each boxplots are total number of biallelic SNPs detected in the population.

For *Pst* isolates in China, the ADMIXTURE showed regional differentiation when K ≧4 (S1 Fig). The *Pst* populations from HB, SC, SX, and GS had similar ancestry, while AH YN, GZ, and CQ had similar ancestral components. In general, the more number (>4) of clusters assigned the more complex ancestry composition was revealed by ADMIXTURE for Chinese *Pst* isolates, while the ancestry remained relatively stable for isolates from non-China.

We next attempted to show how the *Pst* populations were correlated between epidemiological regions and genetic clustering (Fig 2b). The cluster showed the relevance of the epidemic region in China. To explore the genetic composition of the populations from each epidemiological region, and province, we plotted a Sankey plot. As shown in Fig 2b, G2 was detected in most provinces in China but mainly in AutS and SprE regions. G3, G5, and G7 were mainly distributed in SprE, SprS, and AutS regions, respectively. G6 population was detected in SprE, and SprS, but not in AutS. In contrast, *Pst* populations from AutS region were grouped into G2, G3, and G7, while SprS was grouped mainly into G5 and G6, suggesting the divergence and differences in the genetic composition of these two epidemiological regions. It is worth noticing that the *Pst* population from Chongqing (belonging to SprS) was grouped into four genetic clusters. Interestingly, while both Anhui and Hubei belong to the SprE region, the isolates from Anhui had similar ancestry to isolates from SprS (e.g., Yunnan and Guizhou provinces), while isolates from Hubei were more similar in ancestry to isolates from AutS region (e.g., Gansu province), suggesting the different initial inocula in different SprS regions.

We calculated the nucleotide diversity (θ_π_) of each ADMIXTURE-defined population (Fig 2c). The average θ_π_ranged from 0.0035 to 0.0063. The two *Pst* populations from China (G3 and G5) and one international population (G4) had the lower θ_π_ comparing to the remaining populations. In fact, these three populations had the lowest number of biallelic SNPs (Fig 2c). And this low level of nucleotide diversity in the G3, G4, and G5 populations was observed throughout the entire genome (Fig 2c). The alignment results did not show biased sequencing coverages or mapping quality for the isolates in these populations (S1 and S2 Tables). We speculate that the low nucleotide diversity of these three populations might due to the relatively small long term effective population size (See “Demographic history of *Pst* populations”) and strong bottleneck effects. For instance, the isolates in G3 populations (mainly from Hubei province) could not over-summer and must receive large amount of inocula for autumn and spring epidemics every year.

The average Tajima’s *D* values calculated using 10 kb genomic windows along the genome were all positive but varied among *Pst* populations. The international *Pst* group G4 had the lowest average Tajima’s *D* value, 0.5940, while the other international group G1 had the highest value, 3.8823 (Fig 2d). Similar to the case of nucleotide diversity, the low level of Tajima’s *D* values in G4 was also observed throughout the entire genome (Fig 2d). The difference in Tajima’s *D* values suggests that selection forces were imposed on the *Pst* genome in different populations.

Since many alternate host species and the potential for sexual reproduction have been identified in China, we estimated and compared the linkage disequilibrium between the Chinese *Pst* population and the international population excluding China. As expected, the LD coefficients (*R^2^*) decreased as the physical distances between sites increased and had similar decay patterns for both Chinese and international *Pst* populations (Fig 3). For comparison, we also calculated *R^2^* for our previously established *Pst* selfing population. We found that there is a much slower LD decay for the selfing population. As a result, the *R^2^* in the selfing population remained much higher than that of both Chinese and international populations. The slower LD decay was expected since only one cycle of sexual reproduction occurred in the selfing population. Furthermore, we retrieved a natural population of *Puccinia graminis* f. sp. *tritici* (*Pgt*), the causal fungus of wheat stem rust disease, with 77 isolates, and estimated the *R^2^*. The results showed that the *Pgt* had much quicker rate of LD decay (*R^2^* reduced to 0.25 within <1kb) than that in *Pst* populations (*R^2^* reduced to 0.5 within >5kb) (Fig 3). Taking together, these suggested that *Pst* populations have had sexual reproduction in nature, but the frequency were much lower than that in its sister species *Pgt*.

**Fig 3.**
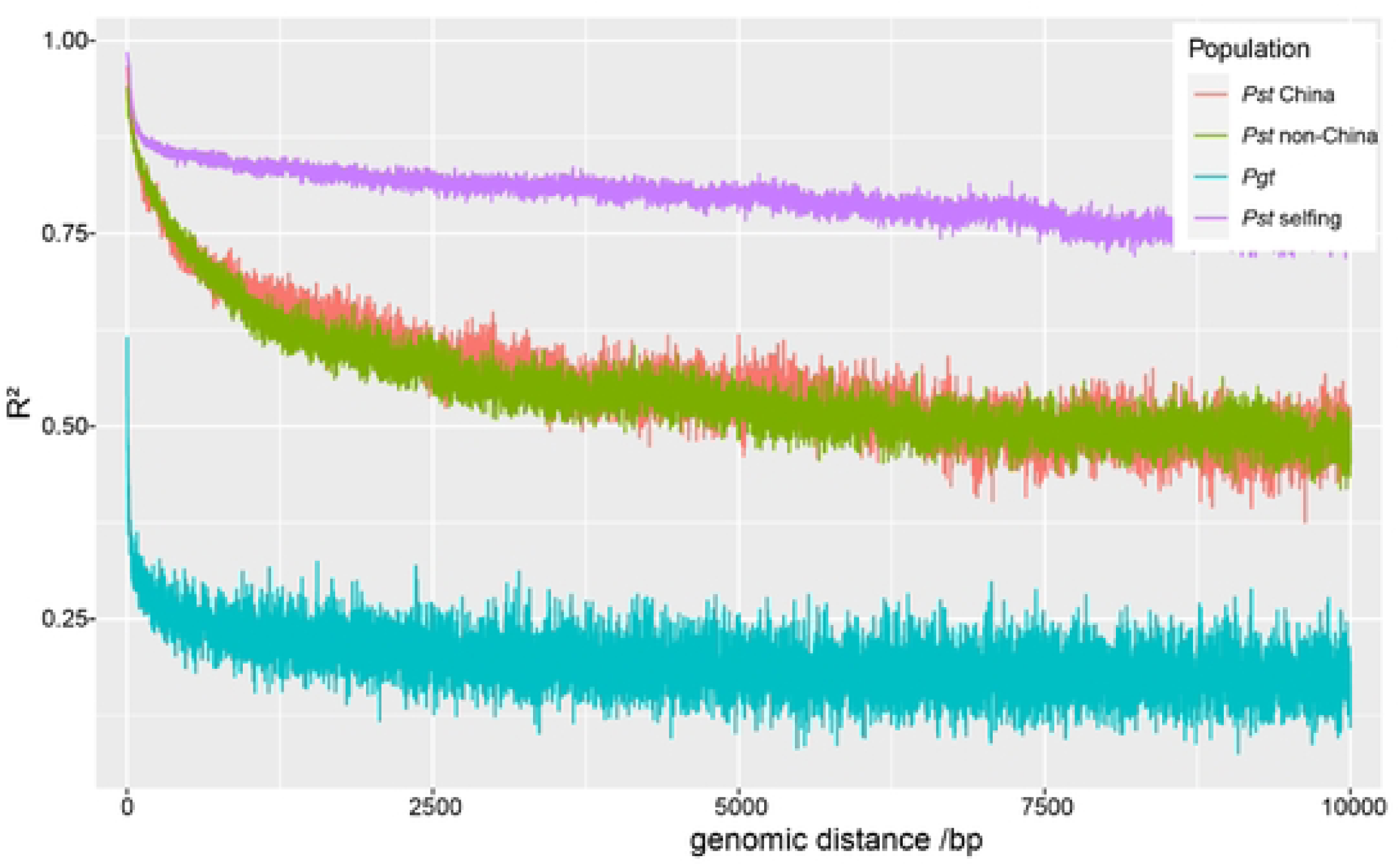
Linkage decay of the *Pst* populations and comparison to *Puccinia graminis* f. sp. *tritici* (*Pgt*), the causal fungus of stem rust in wheat. Red lines, the *Pst* population in China; green line, the international *Pst* population excluding Chinese isolates; blue line, the selfing population; green line, the *Pgt* population with 77 isolates from Guo et al. (2022).

### Genome scan of selective sweeps in Chinese *Pst* populations

The 99.5 percentiles of the CLR distributions built using the highest values from each simulated dataset were from 3 to 22 (Fig. S2), which is much lower than the 99.5 percentiles of CLRs from real populations (ranged from 50.33 to 286.15). The similar situation was also reported in the *Pgt* populations (Guo et al., 2022). Therefore, the empirically defined thresholds of 99.5% were more conservative and should reduce the rate of false positives in the selection analyses. Using such a threshold, 23 to 97 regions harboring hard sweeps were identified in five ADMIXTURE defined Chinese *Pst* populations (Table 1). These regions were from all 18 chromosomes covering 3.98% of the whole genome (Fig 4). The length of single regions ranged from 5.0 Kb to 76.8 Kb.

**Fig 4.**
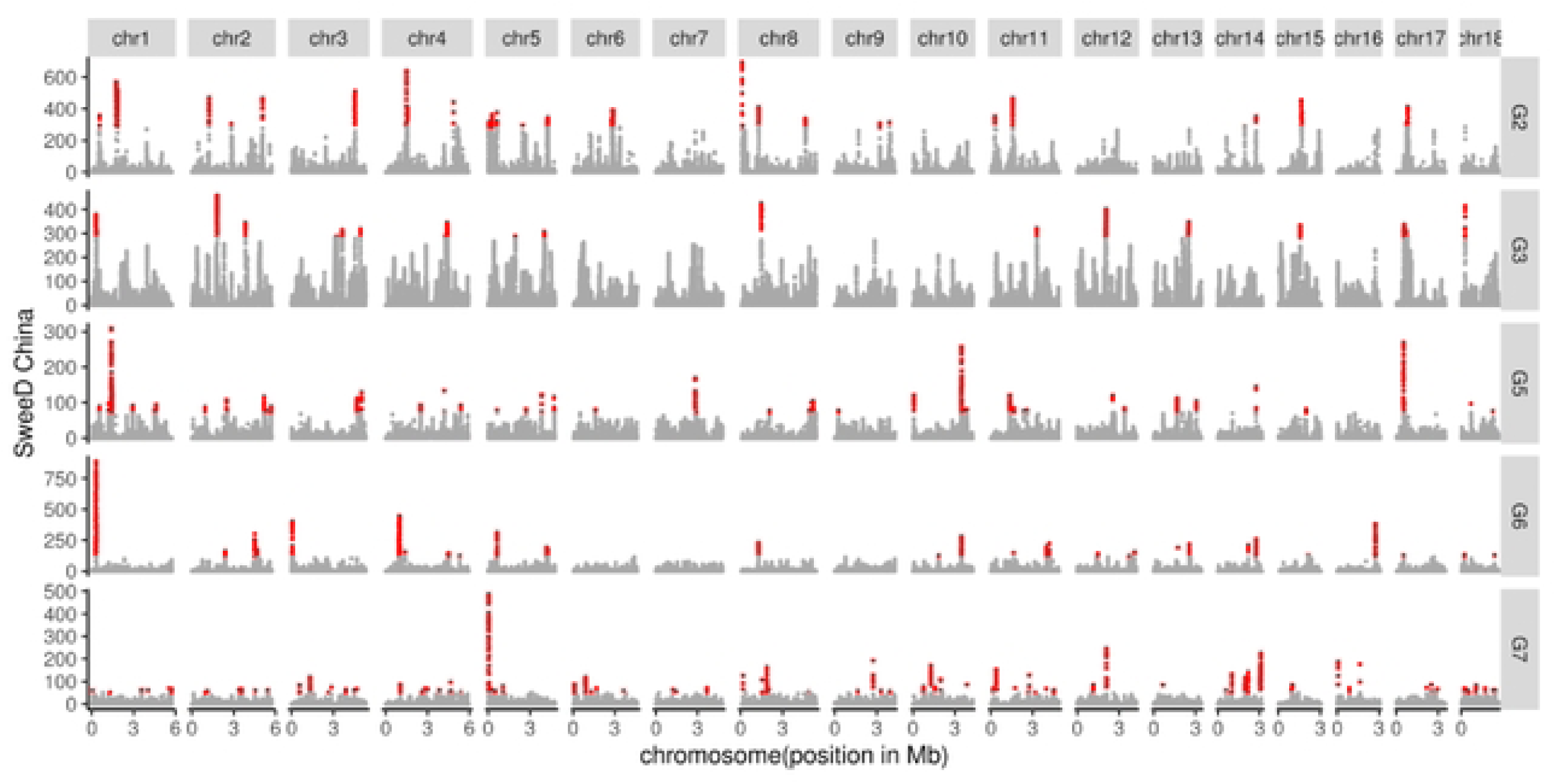
Genome-wide hard sweeps in five Chinese *Pst* populations detected by CLR. Each column represents a chromosome. Each panel represents a group and red dots are outlier sites over 99.5% threshold.

**Table 1.** Summary statistics of the selective sweep regions identified in Chinese *Pst* populations.

| Detection method | Population | Number of selective sweep regions | Total length of regions affected by sweep |  | Number of genes in all sweep regions |
| --- | --- | --- | --- | --- | --- |
|  |  |  | Accumulative | % of whole |  |
|  |  |  | size Kb | genome |  |
| **CLR** | G2 | 44 | 581.9 | 0.74 | 112 |
|  | G3 | 23 | 504.4 | 0.64 | 100 |
|  | G5 | 56 | 676.7 | 0.86 | 195 |
|  | G6 | 35 | 572.9 | 0.72 | 131 |
|  | G7 | 97 | 808.8 | 1.02 | 183 |
| **iHS** | G2 | 209 | 1196.3 | 1.51 | 376 |
|  | G3 | 158 | 870.9 | 1.10 | 271 |
|  | G5 | 57 | 320.3 | 0.40 | 106 |
|  | G6 | 61 | 367.1 | 0.46 | 120 |
|  | G7 | 65 | 371.4 | 0.47 | 115 |

We also performed integrated haplotype score (iHS) statistics to detect soft sweeps using a 99.5% threshold. In G2, G3, G5, G6, and G7, a total of 209, 158, 57, 61, and 65 selective sweep regions were detected as having undergone recent positive selection, covering 3.95% of the genome (Fig 5). The length of individual regions ranged from 10.0 Kb to 21.7Kb.

**Fig 5.**
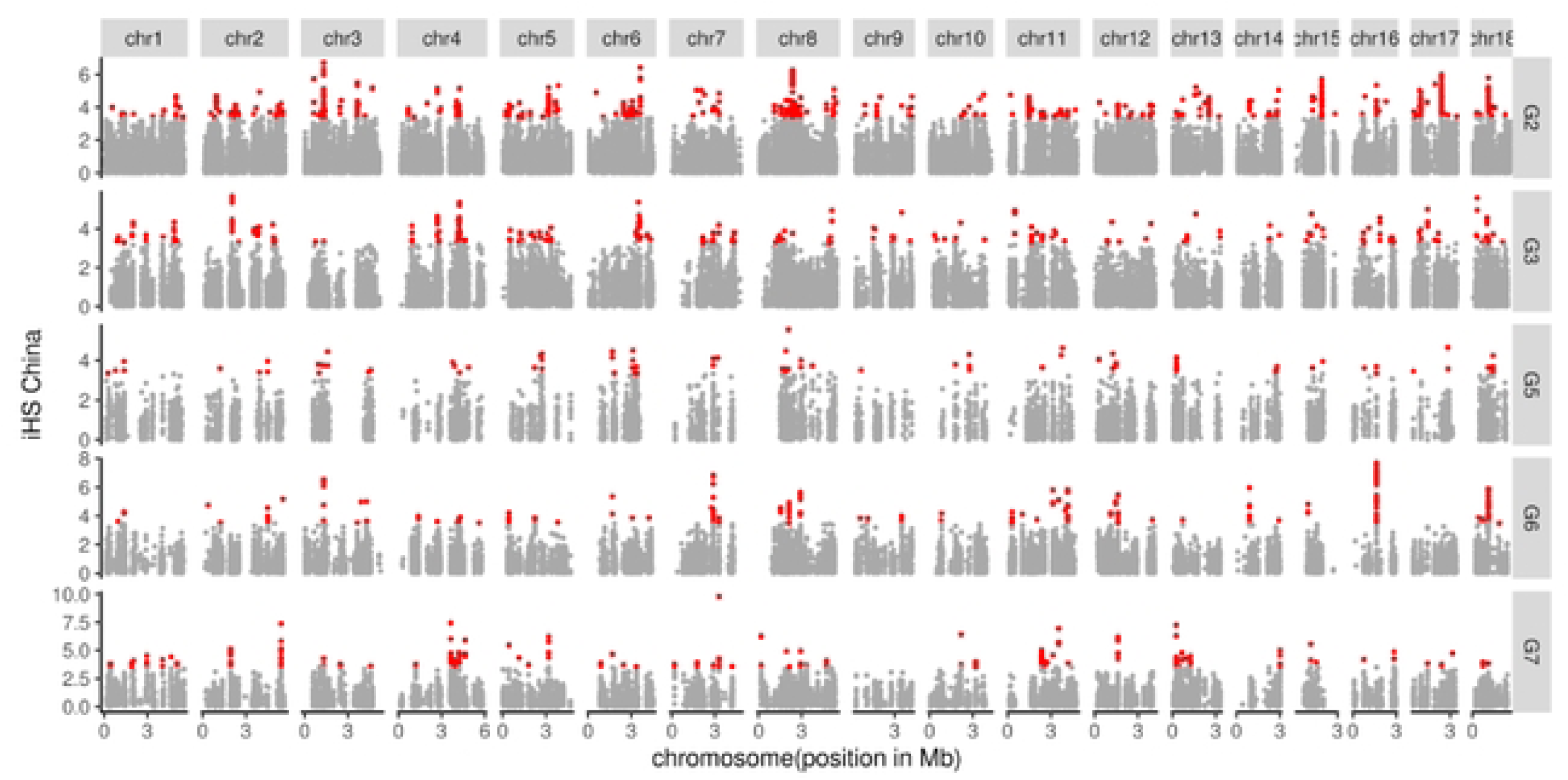
Genome-wide soft sweeps in five Chinese *Pst* populations detected by iHS. Each column represents a chromosome. Each panel represents a group and red dots correspond to outlier sites over 99.5% threshold.

### Gene Function in Selective Sweep Regions

After annotation, we detected 1526 and 988 genes in selective sweep regions detected by CLR and iHS, respectively. Among them, 11.4% and 10.5% were predicted to be secreted protein (SP) coding genes, which was higher than the genome-wide average (9.7%) (Table 2). Further, 33.3% and 33.6% of the SP genes in selective sweep regions were predicted to be effector genes, which is again higher than the genome-wide average (32.6%). A large number of the genes within sweeps were predicted to have reduced virulence when mutated, suggesting they are involved in pathogen-host interactions (S3 Table).

**Table 2.** Content of genes, predicted secreted proteins and predicted effectors in the whole genome and selective sweep regions detected by CLR, and iHS.

| **Region** | **Detection Method** | **Number of genes** | **Gene density (per 10Kb)<sup>a</sup>** | **Number of secreted proteins** | **Secreted protein ratio (%)<sup>b</sup>** | **Number of effectors** | **Effector ratio (%)<sup>c</sup>** |
| --- | --- | --- | --- | --- | --- | --- | --- |
| **Selective sweep regions** | CLR | 1526 | 2.36 | 174 | 11.40 | 58 | 33.3 |
|  | iHS | 988 | 3.16 | 104 | 10.52 | 35 | 33.6 |
| **Genome** |  | 16455 | 2.1 | 1602 | 9.7 | 523 | 32.6 |

<sup>a</sup> Gene density was calculated by dividing the number of genes by the length of total regions.
<sup>b</sup> Secreted protein ratio was calculated by dividing the number of secreted proteins by the number of genes.
<sup>c</sup> Effector ratio was calculated by dividing the number of effectors by the number of secreted proteins.

GO enrichment analysis showed that the genes within soft sweeps were enriched in chitin synthase activity (GO:0004100), hexosyltransferase activity (GO:0016758), ribosome (GO:0005840), and other functions (Table 3). We also found the soft sweeps were enriched in binding activities, such as 48 genes associated to protein binding activities (GO:0005515), 32 genes associated to nucleic acid binding (GO:0003676), 25 genes to zinc ion binding (GO:0008270), 21 genes to DNA binding (GO:0003677) (S4 Table; Fig. S3). Genes within iHS selective sweep had mRNA-related functions, such as translation initiation (GO:0006413), mRNA splicing via spliceosome (GO:0000398), and regulation of transcription by RNA polymerase II (GO:0006357). Similar to gene functions in the soft sweeps, the hard sweeps also had a large number of genes associated with binding activities (Table 3; S5 Table; Fig. S4).

**Table 3.** Gene Ontology functional enrichment of genes in selective sweep regions detected by CLR and iHS (pvalue<0.05)

| Methods | GO ID | GO description | pvalue | ontology<sup>a</sup> |
| --- | --- | --- | --- | --- |
| **CLR** | GO:0004100 | Chitin synthase activity | 1.78E-02 | MF |
|  | GO:0006412 | translation | 3.01E-02 | BP |
|  | GO:0003735 | structural constituent of ribosome | 4.46E-02 | BP |
|  | GO:0004222 | metalloendopeptidase activity | 4.93E-02 | BP |
|  | GO:0016758 | hexosyltransferase activity | 1.38E-02 | CC |
|  | GO:0005840 | ribosome | 4.32E-03 | CC |
|  | GO:0000160 | phosphorelay signal transduction system | 2.07E-02 | MF |
|  | GO:0046982 | protein heterodimerization activity | 2.98E-02 | MF |
| **iHS** | GO:0000398 | mRNA splicing, via spliceosome | 2.25E-03 | BP |
|  | GO:0016226 | iron-sulfur cluster assembly | 2.30E-02 | BP |
|  | GO:0005839 | proteasome core complex | 4.22E-02 | BP |
|  | GO:0006357 | regulation of transcription by RNA polymerase II | 4.42E-02 | BP |
|  | GO:0046873 | metal ion transmembrane transporter activity | 1.28E-03 | MF |
|  | GO:0030001 | metal ion transport | 1.93E-03 | MF |
|  | GO:0006413 | translational initiation | 8.62E-03 | MF |
|  | GO:0051603 | proteolysis involved in protein catabolic process | 4.22E-02 | MF |
|  | GO:0004185 | serine-type carboxypeptidase activity | 5.00E-02 | MF |

<sup>a</sup> Gene ontology terms: BP, Biological Processes; CC, Biological Processes; MF, Cellular Components.

In addition, we found several known function effectors were located in the selective sweep regions, such as *PNPi* (AOS49611.1), *PsKPP4*(AXL48515.1), Ras-like C3 botulinum toxin substrate 1 (KNE95594.1) and PGT-hesp-767 (XP_003322973.1) in the soft sweeps and Ras-like protein Rab7 (KNF04400.1) in the hard sweeps.

### Demographic history of *Pst* populations

Lastly, we attempted to infer the historical effective population size of different *Pst* populations. All isolates in S1 and S2 Tables were combined to represent the international *Pst* population (G_*Pst*), and isolates in S1 Table were used to represent the Chinese *Pst* population (G_China). In Fig 6a, we show the inferred size history over the period from 20,000 years to 200 years ago. In the period from 20,000 to 13,000 years ago, the population sizes of G_China and G_*Pst* experienced a highly similar dynamic and remained stable. The size of both populations had a steep decline from 13,000 to 10,000 years ago (Fig 6a), following with a short period of stable. Then the two populations started to diverge, followed by a round of size expansion and reduction from 3,000 to 60 years ago. The two nadirs of effective population sizes suggested that the pathogen populations are experiencing bottleneck events.

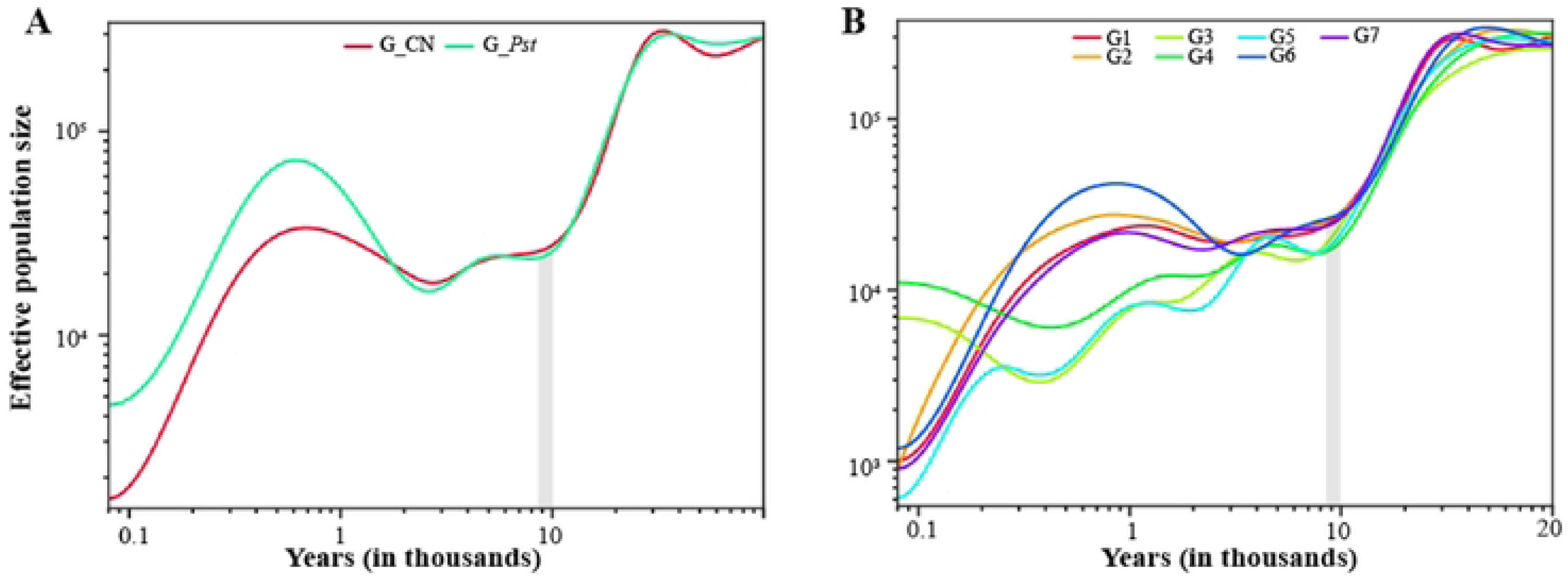

To reveal more details of *Pst* demography, we further inferred the size histories for each ADMIXUTRE-defined population. As shown in Fig 6b, all populations experienced similar dynamics as G_*Pst* and G_China in the distant past, with a steep decline starting 30,000 years ago. The populations diverged starting from 10,000 years ago, and the sizes of populations had a slower decline until 3,000 years ago. Starting from 2,000-3,000 years ago the G1, G2, G6 and G7 populations began to deviate from the rest of the three subpopulations, with a slight increase in population size until 800 years ago followed by a steep decline up to the very recent. Interestingly, the inferred demographic histories of G3 (mainly from Hubei, China), G4 (from India and Ethiopia), and G5 (from Yunnan, China) had similar dynamics, with a moderate decline in size from the divergence with other subpopulations at 300-400 years ago. Lastly, we noted that most *Pst* populations have been experiencing a size decline from recent 100 years to the recent, which might reflect the directional selection of the pathogen by reducing the effective population size from the modern breeding programs.

## Discussion

### Limited gene flow between *Pst* population from China and other countries and low level of sexual reproduction of *Pst* nature populations

Our population genomic analyses unraveled several features of the worldwide *Pst* population. Firstly, Chinese *Pst* populations had limited admixture with that from other countries, suggesting the pathogen might have evolved independently in China considering that the large geographical area and diverse cropping systems provide both over-wintering and over-summering places for *Pst* (Stubbs, 1985; Chen et al., 2013). Secondly, the international *Pst* populations had similar high levels of diversity compared to Chinese populations. Previous studies based on tens of microsatellite markers suggested that populations from Europe, Australia, eastern Africa, India, and the USA were diversity-low while populations from China were diversity-high (Ali et al., 2014). We speculate this inconsistency might stem from the technical bias of genotyping techniques since most microsatellite markers were initially developed and optimized for specific populations but the genome-wide SNPs could minimize such bias. Thirdly, both Chinese and international populations had potential sexual reproduction. Even though the sexual life cycle of *Pst* was discovered a decade ago (Jin et al., 2010), the extent of sexual reproduction in nature still remains unclear. Previous analyses based on several molecular markers could only infer the sexuality of the loci from which the markers were developed. For example, Ali *et al*. (2014) using 20 microsatellites demonstrated that the population of *Pst* on a continent-scale was clonal except the Himalayan and Near-Himalayan (China) regions. To our knowledge, this is the first study using genome-wide loci to infer the extent of linkage disequilibrium in *Pst*. Moreover, we took the advantage of an established selfing population with theoretically one meiosis, as a control for comparison with the natural population. On one hand, the LD decay in both Chinese and international *Pst* populations was much faster than that in the selfing population, suggesting that natural *Pst* had accumulated extensive recombination. On the other hand, the LD decay of *Pst* detected in the present study was slower than that in its sister species *P. graminis* f. sp. *tritici* which has typical sexual reproduction (Fig 3) and other fungi (e.g., *Serpula lacrymans*; Skrede et al., 2021). This confirmed our previous speculation that the frequency of *Pst* sexual reproduction is low in nature. The low frequency of sexual reproduction implies that the importance of alternate hosts in *Pst* epidemics should not be over-estimated.

Historically, three main *Pst* epidemiological regions were designated in China, namely the spring sources of inoculum, autumn sources of inoculum, and spring epidemic areas (Chen et al., 2013). We did not assign clear correspondences between epidemic areas or provinces and genetic populations. This might be explained by the high mixture of ancestry of each isolate which was possibly caused by migration among epidemiological regions/provinces since the germination rate of teliospore is low (Chen et al., 2021) and the frequency of sexual reproduction is rare in nature in China (Wang et al., 2016), which could be support by our linkage disequilibrium analysis (Fig 3). As in autumn sources of inoculum, Gansu and neighboring areas were supposed to be the over-summering region and the origin area of Chinese *Pst* with high diversity and recombination rate (Duan et al., 2010). Traditional epidemiology considered that *Pst* isolates in Gansu had high polymorphism while we found isolates from autumn sources that were mainly grouped to G2 and G7. In contrast to autumn sources of inoculum, spring sources mainly contained G5 and G6 populations. Spring epidemic areas comprised all five populations reflecting the impact of both SprS and AutS. Recently, Li *et al*. (2021) proposed that Yunnan was another origin of Chinese *Pst*, as Yunnan is close to the Himalayans in geography. In contrast to Li *et al*. (2021), we detected the lowest nucleotide diversity in G5 (mostly distributed in Yunnan) plus high Fst between G5 and G1, G4, suggesting the high differentiation from the other countries’ isolates.

### Extensive hard and soft sweeps associated with *Pst* adaptation

Our results indicated that the *Pst* adaptation is associated with both hard and soft selective sweeps. The genome-wide selective signals in *Pst* were 3.95% and 3.98% of the genome. This pervasive selection were also detected in other agricultural plant pathogens (Badouin et al., 2017; Hartmann et al., 2018; Mohd-Assaad et al., 2018), e.g. 0.5% to 4% selective sweep regions in another wheat fungal pathogen *Zymoseptoria tritici*. The classical model of positive selection suggests that the time to adaptation is proportional to the product of the mutation rate of the adaptive variant and the effective population size (Gladieux et al., 2015). Based on this, given the mutation rate of 10^-8^ per site per generation and the effective population size of 10^5^, the time to adaptation for *Pst* would be at the order of thousand generations, which is much slower than the observed (i.e. emergence of new races within only years to decades). One explanation for this conflict is that the assumption that the adaptation is involving a single adaptive variant (that is, hard selective sweep) is not valid in the classical model. Gladieux et al. 2015 proposed that adaptation may be faster given larger mutational targets (i.e. large number of base pairs within an allele, that is, soft selective sweeps). Indeed, our results of balanced hard and soft sweeps in *Pst* are consistent with the soft sweeps-associated rapid adaptation.

Effectors play a crucial part in host-pathogen interactions which makes it undergo selection forces by hosts. For example, filamentous pathogen *Melampsora lini* showed genetic variation at effector loci *AvrP123* and *AvrP4* due to host adaptation (Barrett et al., 2009). In a study of wheat pathogen *Parastagonospora nodorum*, effectors also displayed differentiation driven by local adaptation (Richards et al., 2019). Hence, we performed effectors predicting in the *Pst* genome. A higher ratio of gene density, secreted protein ratio, and effector ratio was displayed in selective sweep regions than in the whole genome of all populations by each testing method. The result implies that pathogenicity genes might undergo recent positive selection. For example, we performed sequence alignment of function-known *Pst* effectors and detected *PNPi* and *PsKPP4* effectors in selective sweep regions. *PNPi* effector interacts with wheat *NPR1* and reduces induction on wheat during infection (Wang et al., 2016). *PsKPP4* strengthens pathogenicity of *Pst* by playing a part in regulating infection-related morphogenesis (Zhu et al., 2018). For the genes within the sweeps but with unknown functions, it would be worthy to explore what specific adaptive phenotypes are associated with these genes.

In selective sweep genes, we detected genes encoding major facilitator superfamily (MFS) transporters, sugar transporter, ATPase family associated with various cellular activities, and heat shock protein (*Hsp*70) family. MFS transporters played roles in pathogen sensitivity to fungal toxins and fungicides, pathogenicity, and response to oxidative stress (Stergiopoulos et al., 2002; Roohparvar et al., 2007). Hartmann *et al*. (2018) detected abundant transmembrane transporter genes in selective sweep regions of continent-scale *Zymoseptoria tritici*, implying selection forces from the environment and host. DNA repair protein swi10 and rad10 in yeast have homologies to the human ERCC1 gene (Rödel et al., 1992). ERCC1 was related to the repair of DNA damage caused by UV light (Rödel et al., 1997). Heat shock proteins play a part in stress conditions in the fungal system and Hsp70 protein was involved in response to cold stress (Tiwari et al., 2015). Temperature and light response reflected selection forces from abiotic factors on *Pst* populations.

### Co-evolution between *Pst* fungi and wheat domestication

Our demography analysis revealed the changes in the effective population size for *Pst* populations along wheat domestication. The sizes of all *Pst* populations had remained relatively stable before the beginning of the hexaploid wheat formation which is estimated to have occured 9,000-10,000 years ago (Lev-Yadun et al., 2000; Levy & Feldman 2022). Remarkably, strong bottleneck events were estimated for all *Pst* populations during the wheat domestication. The bottleneck events could be explained by the restricted genetic variability during wheat domestication (Venske et al., 2019). Nevertheless, this indicated that the demography of *Pst* was affected by the domestication of its host plant. Such an impact of crop domestication on the pathogen has also been observed in the maize-smut pathosystem (Schweizer et al., 2021). While the trends of population sizes differed, we noticed a steady decline for all *Pst* populations in the recent 200 years. Coincidentally, the breeding programs of wheat started during this time frame, and breeding for resistance to stripe rust has been one of the major priorities for wheat (Biffen 1905; Venske et al., 2019). This coincidence led us to speculate that the intentional breeding and deployment of single/major resistance genes could continuously contribute to the decline of *Pst* effective population sizes, even though such kind of resistance could be easily overcome by evolving virulence of *Pst*. However, whether the deployment of multiple genes and even resistance genes from wheat-wild relatives have impacts on *Pst* population demography needs to be further explored, since the more and diverse resistance gene the less selection pressure exerted on the pathogen populations.

In summary, by population genomic analyses of worldwide populations, we revealed unprecedented features of populations of *Puccinia striiformis* f. sp. *tritici*. There might be limited gene flow between *Pst* populations from China and other countries. In terms of selection forces, we found widespread selective sweep regions across the *Pst* genome. We illustrated that the rapid adaptation of *Pst* might be associated with both hard and soft sweeps. Genes within the selective sweeps showed functions related to pathogen-host interaction and environment adaptation. Our study reinforced the view that the fungal plant pathogens, with mainly clonal reproduction and smaller effective population sizes, can have widespread selective sweeps. More studies involving stripe rust pathogen populations from wheat-wild relatives will be needed to illustrate the details of the population evolution and to evaluate the impact of modern agriculture on changes in *Pst* populations. The comprehensive understanding of the co-evolution between pathogens and host plants will enable us to develop sustainable strategies to control the pace of pathogen evolution in agro-ecosystems.

## Materials and methods

### Isolates collection

We selected 69 *Pst* isolates collected from Anhui, Chongqing, Gansu, Guizhou, Hubei, Qinghai, Shaanxi, Sichuan, and Yunnan provinces, covering the major wheat production areas across China, in the year 2015 (S1 Table). After purification from single urediniums, fresh urediniospores were used to inoculate Mingxian169, a susceptible wheat variety to all Chinese *Pst* races, for multiplication. The inoculation procedure was previously described in (Liu et al., 2010).

To explore the population structure between the Chinese *Pst* populations and the international *Pst* populations, sequencing data of twenty-eight worldwide isolates published on NCBI were also retrieved. These isolates were from Australia, Denmark, Ethiopia, Eritrea, Europe, France, India, the UK and the USA (Hubbard et al., 2015; Cuomo et al., 2017; Kiran et al., 2017; Schwessinger et al., 2018; Li et al., 2019; Schwessinger et al., 2020; van Schalkwyk et al., 2021). All the isolates were collected from wheat. It should be noted the international isolates were sampled between 1978 and 2011, while our Chinese isolates were sampled between 2014 Fall to 2015 Spring. Given that the *Pst* was mostly clonal and there was no population stratification by sampling dates (See RESULTS), the difference of sampling date between international isolates and Chinese isolates should have no impact for our analyses.

### DNA extraction and whole-genome sequencing

Genomic DNA was extracted from urediniospores using the CTAB method (Chen et al., 1993). The quality was checked in a 0.8% agarose gel and a ND-1000 spectrophotometer (Bio-Rad, Hercules, CA, USA). Only high-quality genomic DNA with A260/280 ratio between 1.8-2.0 and no contaminating substances was used as input material for the library preparation. Whole genome sequencing was conducted using the Illumina NovaSeq 6000 platform with a 150bp paired-end module in Sinobiocore Biotechnology Co., Ltd.

### Reads processing and genomic variant detection

The sequencing supplier provided raw reads and clean reads for each isolate. The clean reads were generated from raw reads by discarding contaminations or low quality reads when: a) either one read contains adapter contamination; b) more than 10% of bases are “N” in either one read; and c) the proportion of low quality (Phred quality <5) bases is over 50% in either one read. The Illumina clean reads of each isolate were further trimmed, by removing adapter and primer sequences at read ends using Trimmomatic-0.39 (Bolger et al., 2014) with the following settings: LEADING:20 TRAILING:20 SLIDINGWINDOW:4:20 MINLEN:100. The high-quality trimmed reads were mapped to the *Pst* CYR34 reference genome (Xia et al., 2022) using the BWA-MEM (Li & Durbin, 2010) with default parameters except for: -r 1.0. The CYR34 genome was selected as the reference because a) it is the most prevalent race in China, and b) it is well-annotated with high quality (BUSCO completeness of 97.0%). After removing the duplicates (the identical reads mapped the same genomic region, usually generated from library amplification and could result in false positive in variant calling) using SAMtools v0.1.19 (Li 2011), the SAM formatted alignments were converted to the BAM format. The Picard Tools software (http://picard.sourceforge.net/) was used to clean and sort of the BAM files, which were then indexed using SAMtools. To obtain a preliminary quality control before variant calling, qualimap v2.2.1 (Garcia-Alcalde et al., 2012) was used to check the mapping coverage and quality of all isolates.

In order to reduce the false positive and mismatch variants in the alignments, we conducted the GATK haplotypecaller twice following the recommended procedures using the GATK 4.1 (DePristo et al., 2011). First, we performed three steps to obtain high-quality known sites. Calling high false positive variants with an initial round of variant calling using GATK HaplotypeCaller. Then we performed VCFtools v0.1.16 (Danecek et al., 2011) to filter biallelic SNPs and InDels with a minimum quality of 1,000. After indexing the VCF file, a Base Quality Score Recalibration (BQSR) step was conducted to recalibrate. Then the high-quality known sites were subject to the second round of variant calling using GATK HaplotypeCaller for each isolate.

GVCF files of different isolates with only bi-alleles and a minimum variant calling quality of 1,000 were acquired using VCFtools. To combine multi-sample GVCFs, we performed GATK ConbineGVCFs and GenotypeGVCFs and obtained a final joint genotyping VCF file containing all variants of all isolates. Then we performed GATK SelectVariants to separate SNPs and InDels. We only kept high genotyping quality SNPs within coverages from 20X to 120X to reduce false positives in repetitive regions. As rare alleles may cause false positive results in the following analyses, we removed variants with minor allele frequency less than 0.05. These filters were implemented using VCFtools with the parameters set as --min-meanDP 20 --max-meanDP 120 -- minQ 1000 --minGQ 40 --maf 0.05 --max-missing 0.8.

### Population structure and summary statistics

We analyzed the population structure of 97 worldwide *Pst* isolates using two methods, Principal component analysis (PCA) and Admixture. To avoid biases caused by linkage disequilibrium (LD), the SNPs used for detecting population structure were filtered using “--thin 1000” option in VCFtools. After making the BED file with PLINK, we used the GCTA (Yang et al., 2011) software to perform principal component analysis (PCA) on all *Pst* isolates. We set the PC number to 10 and selected the top three principal components to draw a scatter plot with R package ggplot2. Admixture-v1.3.0 (Alexander et al., 2009) software was used to analyze the relationship of all the isolates with K value setting from 2 to 10. For each group, we performed 50 iterations to compare the cross validation (CV) errors of each K value. The K value with the lowest CV value is set as the most suitable population number. The isolate was assigned to specific cluster when it had at least 70% ancestry from that cluster. Otherwise, the isolate was considered as an admixture. The admixed isolates were excluded from subsequent demographic estimates and selective sweeps analyses. To explore the genetic composition of the populations from each epidemiological region, and province, we plotted a Sankey plot, a type of data visualization showing the flow of information (e.g. number of isolates) between different variables (e.g. epidemiological regions, provinces, and genetic clusters).

Genetic differentiation index (*Fst)* and nucleotide diversity (θ_π_) for each of the seven populations across 10 kb windows of the genome were calculated using VCFtools. To detect the departure from the standard neutral model, we calculated Tajima’s *D* statistics (Tajima, 1989) at both the genome-wide level and genomic windows along each chromosome using VCFtools. A Tajima’s *D* value of 0 suggests that the population evolves at mutation and genetic drift equilibrium and that no selection at a specific locus. Positive values of Tajima’s *D* (with low levels of both low and high-frequency polymorphisms) indicate a decrease in population size or balancing selection, while negative values (with excess of low-frequency polymorphisms) suggest population size expansion or directional selection.

We calculated linkage disequilibrium (LD) using VCFtools with --geno-r2, and --ld-window-bp 20000 options, for the Chinese *Pst* population and the international *Pst* population excluding China. The rate of LD decay could indicate the level of sexual reproduction. For comparison, we also re-analyzed our previously established *Pst* selfing population with 93 isolates (Xia et al., 2020). In addition, we retrieved a recently available whole-genome dataset of *Puccinia graminisi* f. sp. *tritici* (*Pgt*) (Guo et al., 2022), the sister species of *Pst* and the casual pathogen of wheat stem rust disease. It is well known that the *Pgt* fungus has frequent sexual reproduction in nature conditions (Leonard & Szabo 2005; Wang et al., 2015). Therefore, it could be used as a reference for evaluation of the extent of sexual reproduction in rust fungi.

### Genome regions scanning for selection

A selective sweep emerges after the frequency of an advantageous genetic variant rises to high. Generally, there are two kinds of selective sweeps depending on the number of advantageous haplotypes in a population. One is hard sweeps in which single selectively advantageous haplotype has high frequency, while there are multiple selectively advantageous haplotypes in the soft sweeps (Garud 2023). To scan selective sweeps in five Chinese *Pst* populations (G2, G3, G5, G6, and G7; see Results), we used two complementary methods, the composite likelihood ratio (CLR) method for hard sweeps and integrated haplotype score (iHS) method for soft sweeps. The CLR is a composite likelihood approach to compare the likelihood of a model of positive selection to a model of neutral evolution, given that the allele frequency spectrum of the population will be different from that expected under neutral evolution if a population is under positive selection (Pavlidis et al., 2013). Before scanning, haplotype phasing was performed using SHAPEIT4 (Delaneau et al., 2019). First, CLR test based on site frequency spectrum (SFS) was implemented by the SweeD (Pavlidis et al., 2013) software, as this method allows the detection of intra-population hard sweeps (Pavlidis & Alachiotis, 2017). This increases the robustness of demographic models because the null model relies on the variation of the sequence SFS along the whole genome instead of the standard neutral model (Nielsen, 2005; Pavlidis et al., 2013). To determine the threshold for selection analysis, we computed the distributions of CLRs in 1000 data sets per population simulated under the inferred demographic models (see Section ‘Demographic reconstruction’) for each population. To reduce computation burden, only one chromosome with 6 Mb was simulated. The time and population size were scaled to 4N_0_ and N_0_, respectively. The growth rate α at time *t* was calculated according to N(*t*) = N_0_*exp^-α*t*^, where N0 is the initial population size. Five to eight timepoints from best demographic history model were used in simulation by setting -eN and -eN parameters in MacS (Chen et al., 2019). The CLRs were generated for each simulated dataset using SweeD. In addition, we searched soft sweeps using integrated haplotype score (iHS) test using the R package REHH2.0 (Gautier et al., 2017). The iHS test compares ancestral alleles Extended Haplotype Homozygosity (EHH) and derived alleles EHH of each SNP to detect lengthy haplotype blocks. We set the maxgap option to 20,000, and limehh to 0.05 while using scan_hh. Outliers within 5Kb were regarded as the same selective sweep area. We obtained the selective sweep region by adding 5 kb at the flanks of each outlier site. As low SNP density can cause false positive in CLR test, we computed SNP density in 50 kb nonoverlapping windows along the whole genome. Only CLR outlier sites located in a window whose density was above 100 were retained. Gene ontology (GO) terms were assigned to the genes within sweeps by searching Pfam, SUPERFAMILY, TIGRFAM, SFLD, CDD, PIRSR, and PANTHER databases. Enrichment was performed using clusterProfiler (v4.6.0; Wu et al., 2021) package in R.

### Demographic reconstruction

We exploited SMC++ (Terhorst et al., 2017) software to construct the demographic history of Chinese *Pst* populations with whole genome sequences. SMC++ combined the SFS and the coalescent hidden Markov Models (HMMs) to calculate changes in effective population size over time, which makes the inference computationally efficient and accurate. The SMC++ is robust to phasing errors and sample sizes, usually 2-10 individuals with high sequencing coverage were recommended in practice (Terhorst et al., 2017). To distinguish missing regions from very long runs of homozygosity, the raw variants were filtered by setting --minGQ 60 using VCFtools (Danecek et al., 2011) and low-quality variants were masked as missing, following SMC++ recommendation (https://github.com/popgenmethods/smcpp). SMC format was converted by the first dataset using smcpp vcf2smc and then used to estimate effective population size with smcpp estimate. For each population, four isolates were randomly selected as distinguished lineages for vcf2smc (specified by -d option). We set the mutation rate as 2e^-8^ per base pair per generation according to Xia et al. (2018). Finally, smcpp plot was used to plot the effective population size over time. To convert coalescent units from generations to years, we considered the epidemiology in Gansu province since *Pst* could over-summer and over-winter in this region and then served as the major center of inoculum sources for Eastern China. In the Gansu province, there is generally one generation before *Pst* over-wintering. Then in the following spring, sporulation starts from March and lasts to June, with one generation per month. Therefore, five generations per year was used in the demographic history inference.

### Gene function annotation and enrichment Analysis

Genes detected in the selective sweep regions were functionally predicted using InterProScan v.5.51-85.0 (Jones et al., 2014). We used a local lookup to assign these genes to protein families (Pfam), gene ontology (GO) terms, SUPERFAMILY, TIGRFAM, SFLD, CDD, PIRSR, and PANTHER. We also predicted signal peptide and transmembrane signatures with SignalP 4.1 (Petersen et al., 2011), and TMHMM 2.0 (Krogh et al., 2001), respectively. Proteins containing a signal peptide predicted in the SignalP were defined as secreted proteins, after removing proteins predicted as transmembrane by TMHMM.

To detect the phenotype of mutants, we aligned secreted proteins to Pathogen-Host Interaction database (PHI-base) version 4.10, released on 2^nd^ November 2020, (Urban et al., 2020) using BLASTP 2.9.0. We performed GO enrichment analyses for all genes located in selective sweep regions detected by CLR, iHS, and XP-EHH, respectively. Enrichment analyses of GO terms were performed using the R package clusterProfiler (Yu et al., 2012) with a false discovery rate (p-value) set to 0.05. Only GO terms with at least five different genes assigned in the reference genome were considered. We also searched effector protein sequences of known function in the CYR34 reference genome to detect effectors harboring in the selective sweep regions.

## Acknowledgements

We thank Dr. Yuanwen Guo from Kansas State University for the help on the phylogenomic analyses and valuable discussions.

## Author contributions

TL, WC and CX designed the research; YX performed the research; YX, LH, NZ, HL, XZ, AQ, WT, MW, XC, HZ, BL and TL contributed materials and resources, and analyzed the data; YX and CX wrote the paper. All authors read and approved the final manuscript.

## DATA AVAILABILITY STATEMENT

The raw sequence data reported in this paper have been deposited in the Genome Sequence Archive in National Genomics Data Center, China National Center for Bioinformation / Beijing Institute of Genomics, Chinese Academy of Sciences (GSA: CRA010026) that are publicly accessible at https://ngdc.cncb.ac.cn/gsa.

## Supplementary Figures

**S1 Fig.**
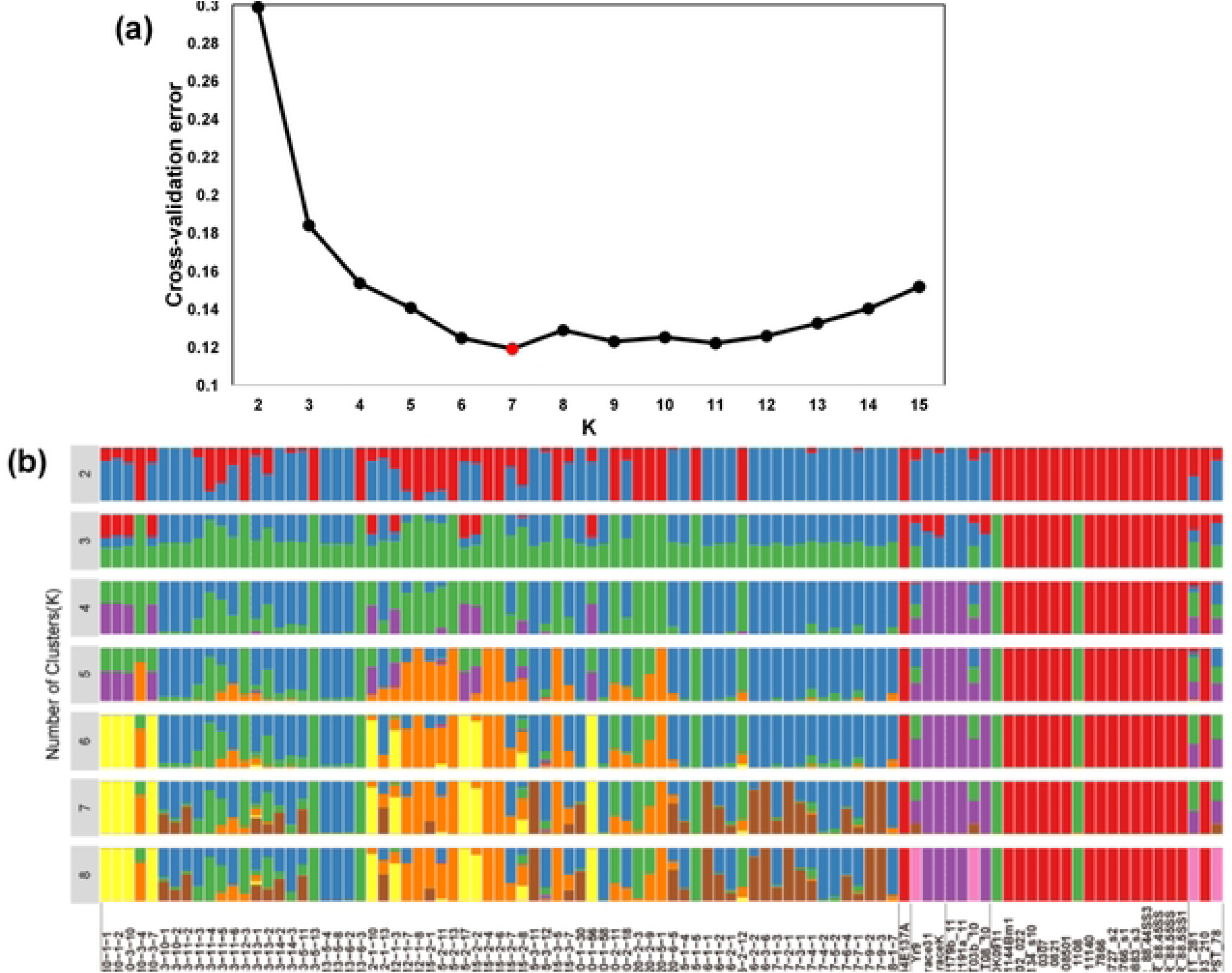
Population structure of 98 worldwide isolates of *Puccinia striiformis* f. sp. *tritici*. A. Plot of cross validation errors of Admixture from K=2 to K=15. K=7 was chosen with the minimized cross validation error. B. Genetic cluster inferred by Admixture software, showing 2 to 8 clusters.

**S2 Fig.**
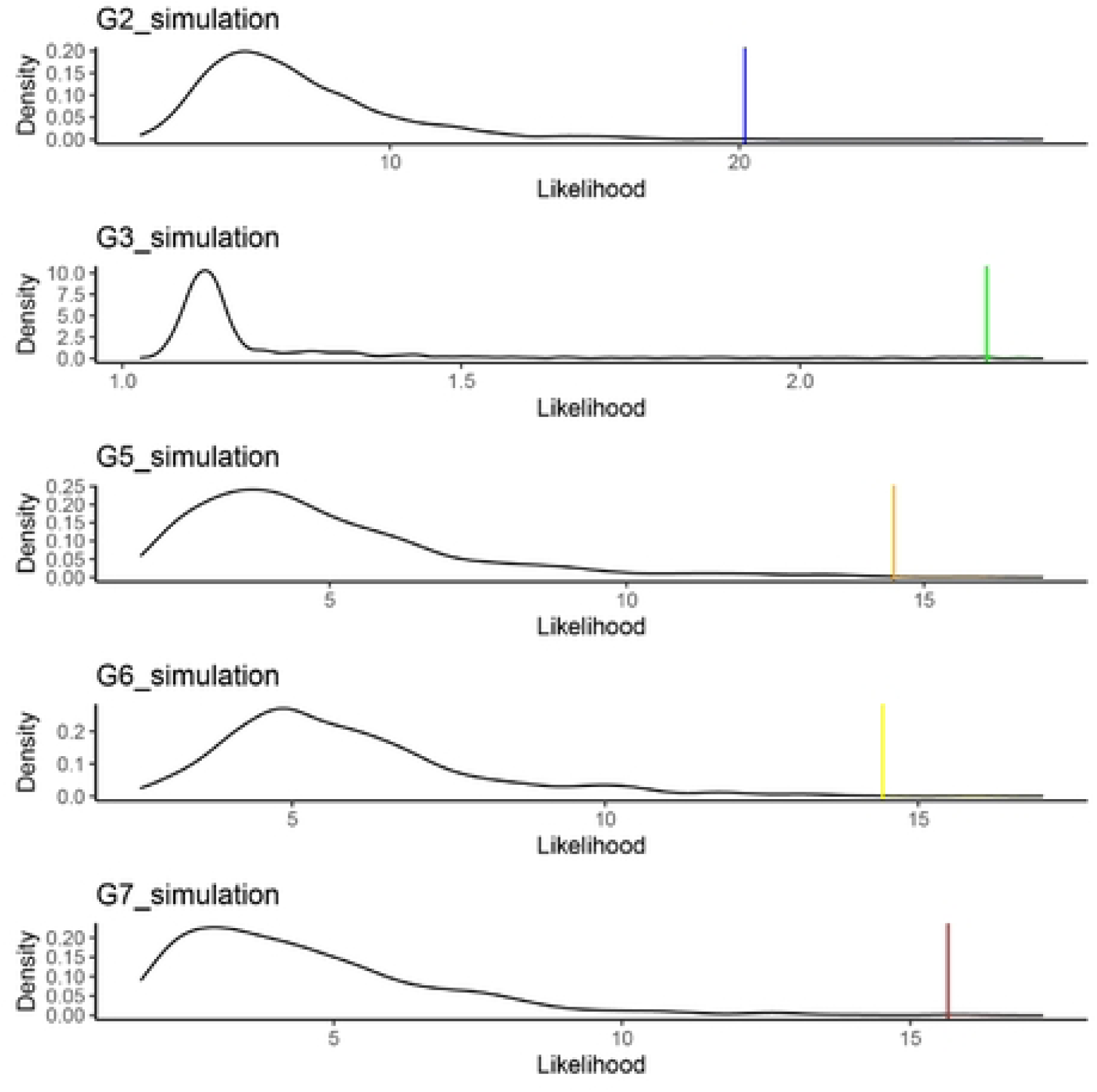
The distribution of composite likelihood ratios (CLRs) simulated under the inferred demographic models for each population. The colored vertical bars at the tail of the distribution indicate the 99.5 percentiles.

**S3 Fig.**
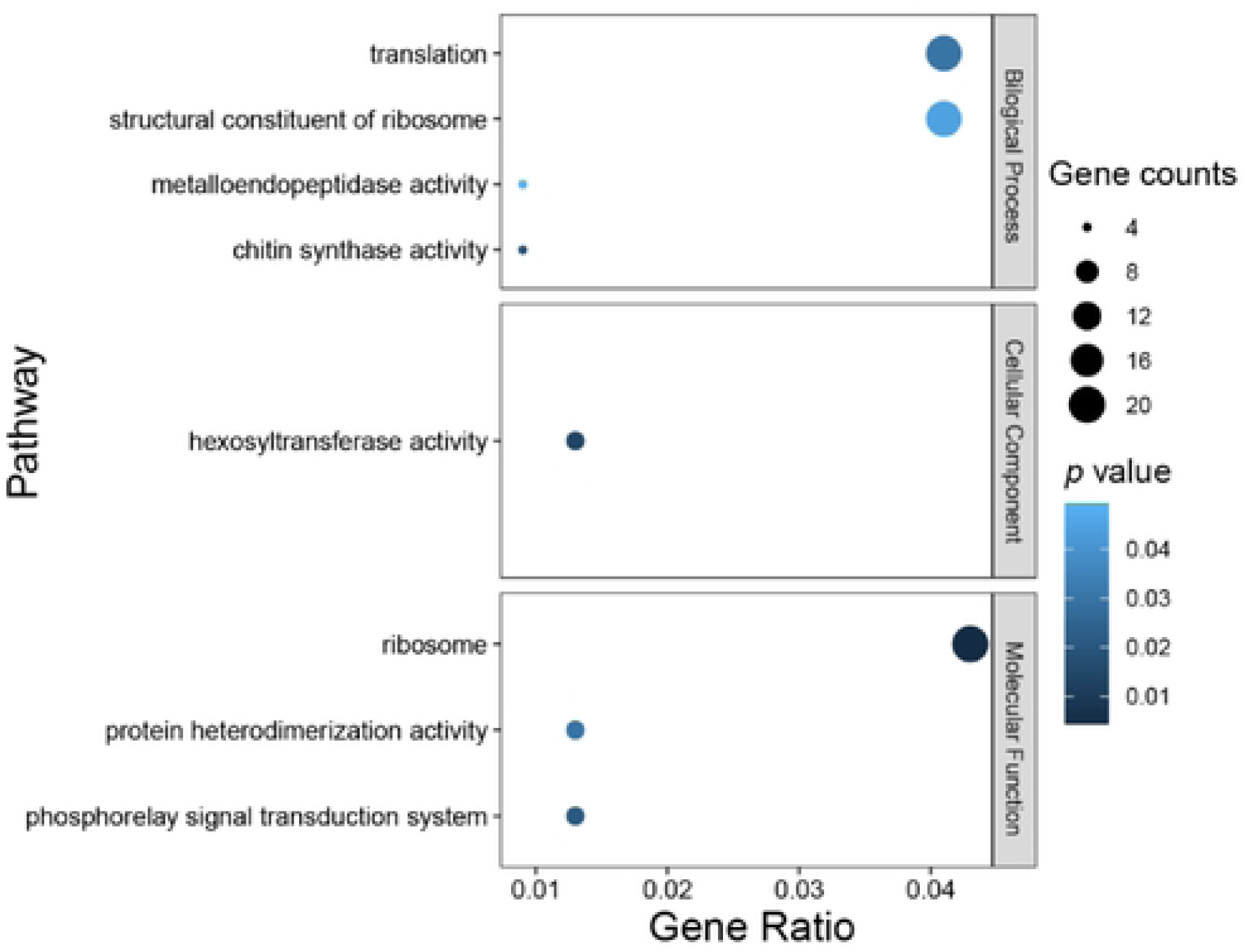
GO enrichment of genes within soft sweeps detected by CLR.

**S4 Fig.**
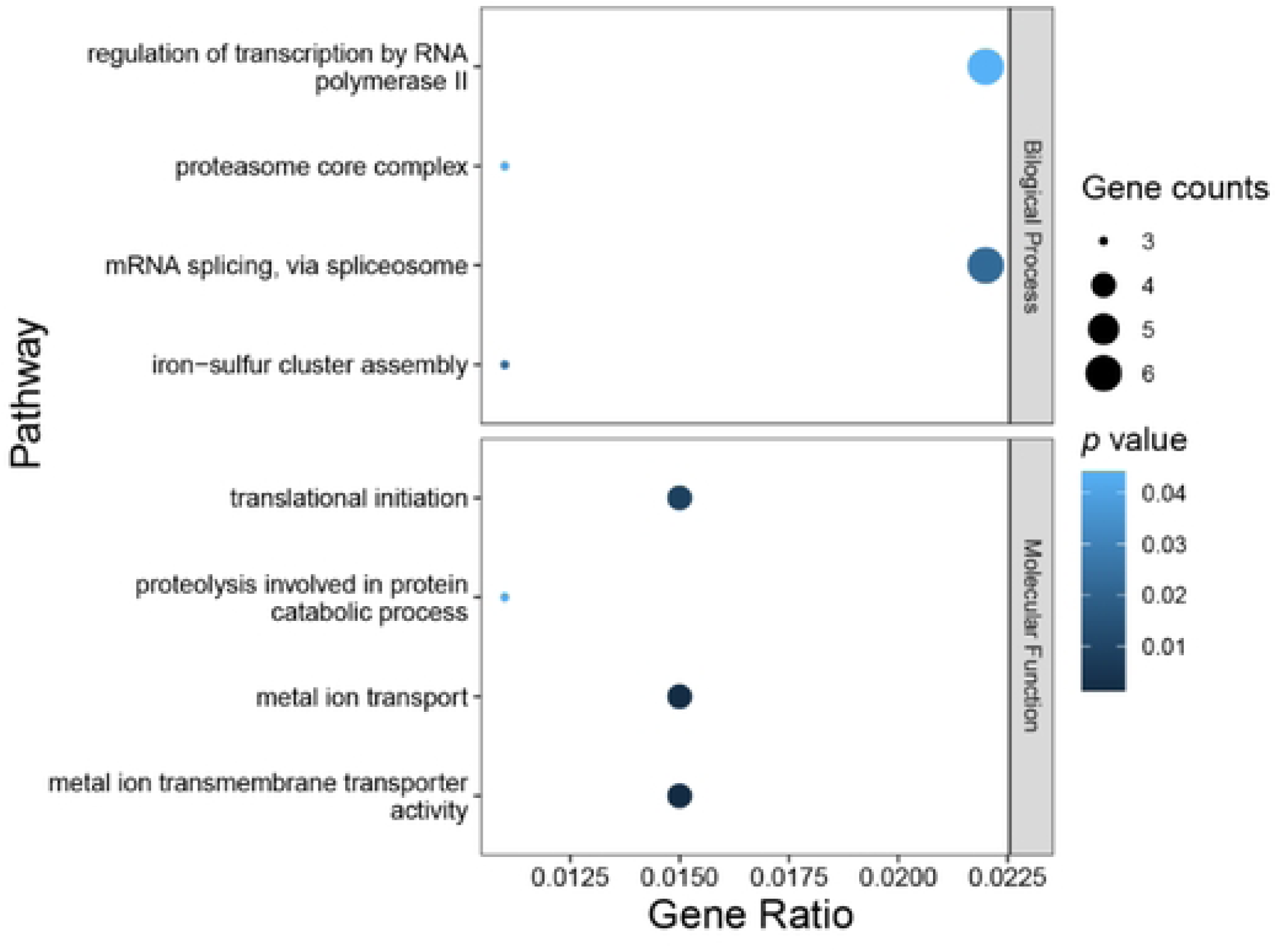
GO enrichment of genes within soft sweeps detected by iHS.

## Supplementary Tables

**S1 Table.** Information of Pst isolate collections from Anhui, Chongqing, Hubei, Gansu, Guizhou, Qinghai, Shaanxi, Sichuan and Yunnan provinces in China

**S2 Table.** Information of Pst isolates from Australian, India, Ethiopia, Eritrea, Denmark, the UK, the USA and France. Raw SRA data of each isolate were downloaded from the NCBI.

**S3 Table.** Mutant phenotype of predicted effectors in selective sweep regions detected by CLR and iHS (evalue<1e-5)

**S4 Table.** GO enrichment of genes within soft sweeps detected by CLR

**S5 Table.** GO enrichment of genes within soft sweeps detected by iHS

